# Decoding individual differences in mental information from human brain response predicted by convolutional neural networks

**DOI:** 10.1101/2022.05.16.492029

**Authors:** Kiichi Kawahata, Jiaxin Wang, Antoine Blanc, Naoya Maeda, Shinji Nishimoto, Satoshi Nishida

## Abstract

Recent advantages in brain decoding with functional magnetic resonance imaging (fMRI) have enabled us to estimate individual differences in mental information from brain responses to natural stimuli. However, the physical constraints and costs of fMRI measurements prevent brain decoding from achieving real-world applications. We address this issue by building a novel framework to decode individual differences in mental information under natural situations from brain responses predicted using convolutional neural networks (CNNs). Once the CNN-based prediction model is trained using measured response, mental information can be decoded from the predicted responses of individual brains with no need for additional fMRI measurements. The model was found to capture individual-difference patterns consistent with conventional decoding using measured responses in 81/87 items to be decoded. Our framework has great potential to decode personal mental information with low dependence on fMRI measurements, which could help substantially expand the applicability of brain decoding in daily life.

## 1. Introduction

Brain decoding based on functional magnetic resonance imaging (fMRI) has been a valuable tool for not only neuroscience research (J. V. Haxby et al., 2014; Tong & Pratte, 2012; van Gerven et al., 2019) but also real-world applications, such as neuromarketing (He et al., 2021; Plassmann et al., 2015). Recent fMRI-based decoding techniques were used to successfully recover rich mental information from fMRI signals induced by natural scenes (Huth et al., 2016; Matsuo et al., 2016; Nishida & Nishimoto, 2018; Nishimoto et al., 2011; Seeliger et al., 2018), and also to attempt to decode the individual differences in mental information, for example, when personal episodes of daily experiences are recalled (Anderson et al., 2020). Although these decoding techniques have capabilities with promising real-world applications, the physical constraints and the high costs of the required fMRI measurements prevent these techniques from achieving widespread application.

One approach to solve this problem uses alternative neuroimaging devices, such as electroencephalography (EEG), that require weaker constraints and lower costs (Rashid et al., 2020; Saeidi et al., 2021). However, signals collected with these devices are noisier and have lower spatial resolution than fMRI, making them less suited to recover rich mental information (Bigdely-Shamlo et al., 2015; Jas et al., 2017) and limiting their real-world application for brain decoding. Another approach uses computational methods that can eliminate a large part of fMRI measurement needed for brain decoding. An example of such a method is the decoding of fMRI responses predicted by computational models (predicted-response decoding) instead of measured responses (Nishida et al., 2020). Previous work has demonstrated that predicted-response decoding achieves the performance comparable to the decoding of measured fMRI responses (measured-response decoding) (Nishida et al., 2020). The predicted-response decoding has the potential to drastically reduce the constraints and the costs of fMRI measurements and expand the applicability of brain decoding. However, the previous work only compared its group-level performance with measured-response decoding, leaving its full potential largely unexplored.

Here we aimed to extend our previous work on predicted-response decoding (Nishida et al., 2020) to the estimation of individual differences in mental information evoked by natural scenes. We have constructed prediction models that simulate the fMRI responses of an individual to arbitrary natural movie scenes and decoding models that estimate an individual’s mental information associated with the viewed scenes from the predicted responses within the whole cortex (Fig. 1). To validate these models, we used a large variety of decoded items associated with natural movie scenes, and evaluated the accuracy with which model estimates captured the individual differences in mental information derived from measured-response decoding. Our results indicate that the framework based on predicted-response decoding has the potential for weak-constraint, low-cost brain decoding to accurately estimate rich mental information varying from person to person.

**Fig. 1.**
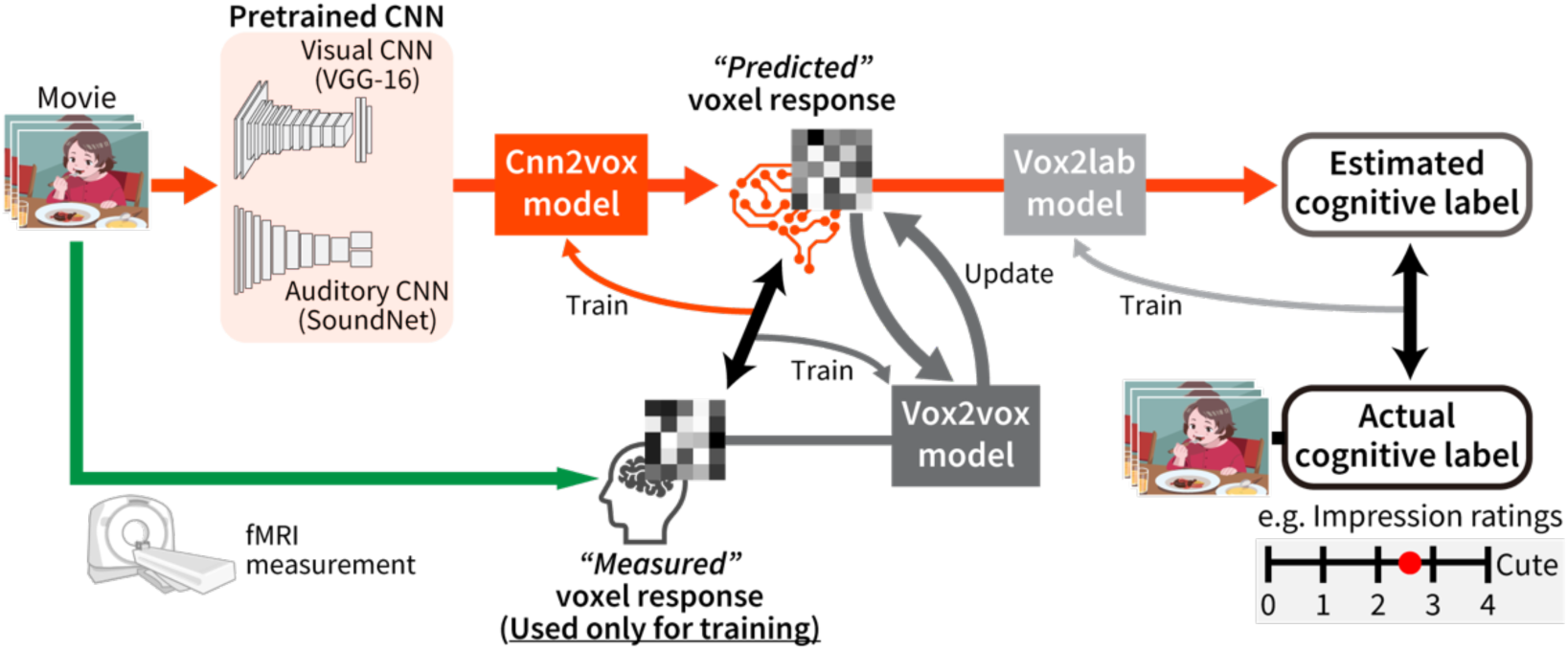
Predicted-response decoding. The framework, applied for each participant, works in two steps: (i) predicting voxel responses to movie scenes, by cnn2vox and vox2vox models, and (ii) decoding the predicted voxel responses to estimate cognitive labels associated with the scenes, by vox2lab models. The models are trained by linear regression using voxel responses measured with fMRI. The cnn2vox model transforms audiovisual features extracted via pretrained CNNs (VGG-16 and SoundNet) into voxel responses. The vox2vox model updates the predicted responses using the history of preceding voxel responses. The vox2lab models are trained by linear regression using paired data of predicted voxel responses and cognitive labels.

## 2. Methods

### 2.1. Predicted-response decoding framework

#### 2.1.1. Overview

Our framework based on predicted-response decoding consists of models to predict fMRI voxel responses to movie scenes (CNN-to-voxel [cnn2vox] and voxel-to-voxel [vox2vox] models) and models to decode movie-associated cognitive labels from the predicted voxel responses (voxel-to-label [vox2lab] models) (Fig. 1). First, the cnn2vox model transforms the features of movie scenes, extracted via visual and auditory CNNs, to voxel responses. Next, the vox2vox model modifies the predicted voxel response using the history of preceding voxel responses. Finally, the vox2lab model estimates cognitive labels from the modified responses.

The cnn2vox and vox2vox models are pretrained on small datasets of movie-evoked responses collected from individual brains using fMRI. The important advantage of this approach is that once the training is completed, these models predict voxel responses to arbitrary movie scenes by transforming the CNN features to the response of individual brains with no need for additional brain measurement. In the last stage, the vox2lab model is trained using paired datasets of predicted voxel response and cognitive labels linked with movie scenes.

#### 2.1.2. CNN feature extraction

VGG-16 (Simonyan & Zisserman, 2014) and SoundNet (Aytar et al., 2016), which were pretrained and available on the web, were used to extract visual and acoustic features, respectively, from movies. To extract visual features from movies via VGG-16 (originally applied to static images of fixed size of 224 × 224 pixels), the movies were decomposed into frames and resized to the same size. Then, unit activations of layers were calculated and pooled for each second of the input stream of the movie frames. Finally, the maximum activation value of each unit for each second was used as the visual features of the movies. We used five pooling layers (pool1–5) and three fully-connected layers (fc6–8) and obtained the visual features for each layer. To extract acoustic features from the same movies, the sound waves of the inputs were resampled at a fixed frequency of 44100 Hz and decomposed into each second. Then, unit activations of layers in SoundNet were calculated for an input stream of sound waves as the acoustic features of the movies. We used seven convolutional layers (conv1–7) and obtained the acoustic features for each layer. Thus, these feature extraction processes produced eight series of visual features and seven series of acoustic features of movies.

#### 2.1.3. Cnn2vox model

The construction of the cnn2vox, vox2vox, and vox2lab models is based on the voxelwise modeling technique (Naselaris et al., 2011). Using a time series of CNN features and voxel responses, the cnn2vox model acquires the linear mapping from a CNN feature space to a response space of each voxel through statistical learning. The learning objective is to estimate the weights of *N* voxels, denoted by W_CV_ = {W_CV(1)_, …, W_CV(*N*)_}, used in the linear model: R = *f*(X)W_CV_ + *∊*, where R = {r_1_, …, r*_N_*} is a series of responses in each of *N* voxels, X is a series of movie scenes, *f*(X) is its feature representation with the dimensionality of *D*, and *ϵ* is isotropic Gaussian noise. A set of linear temporal filters is used to capture the hemodynamic delay in the fMRI response (Nishimoto et al., 2011). The matrix *f*(X) was constructed by concatenating four sets of *D*-dimensional feature vectors with temporal shifts of 3, 4, 5, and 6 seconds. This means that voxel response at a time point *t*, as denoted by R_(*t*)_ = (*t* = 1, …, *T*), was modeled by a weighted linear combination of the preceding series of CNN features:

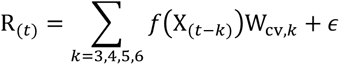

where W_CV,*k*_ denotes the weights corresponding to the delay *k*. The weight estimation was performed using L2-regularized linear least-squares regression. The optimal regularization parameter for each model was determined by 10-fold cross validation of training data and shared across all voxels. In this study, *f*(X) represented a series of unit activations induced by movies for each layer of VGG-16 or SoundNet. Since the substantial number of units in lower layers of the CNNs imposed a large computational cost for the regression process, the dimensionality of unit-activation features, *D*, for each layer was reduced to 1000 in advance by principal component analysis on training data. Finally, eight models for eight VGG-16 layers and seven models for seven SoundNet layers were constructed for each brain.

Each of the estimated linear models predicts voxel responses to new movie scenes. Then, the predicted voxel responses from individual models are integrated using linearly weighted averages for each voxel. The weight for a given voxel is calculated based on the prediction accuracy (Pearson correlation coefficient between measured and predicted voxel responses) for that voxel calculated during the cross-validation in model training. Specifically, the weight for the *i*-th model, denoted by *w_i_*, was determined as 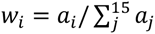, where *a_i_* is the prediction accuracy of the model. This integration process produced a single series of predicted voxel responses to the new movie scenes.

#### 2.1.4. Vox2vox model

The vox2vox model predicts response in one voxel at a given time point from responses in a group of voxels at preceding time points. Hence, this model captures endogenous properties of voxel responses, such as intrinsic connectivity between different brain regions (Fox & Raichle, 2007); in contrast, the cnn2vox model captures exogenous properties of voxel responses, such as stimulus selectivity. The vox2vox model was reported to improve performance in predicting voxel response to movie scenes (Nishida et al., 2020).

In the vox2vox model, the response in each of *N* voxels at a time point *t*, denoted by R_(*t*)_, was modeled by a weighted linear combination of responses in the set of *M* voxels preceded by 1, 2, and 3 seconds:

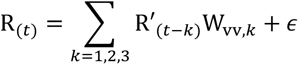

The *M* voxels were selected based on the cnn2vox model prediction accuracy on training data. We used the top 2000 voxels with the highest prediction accuracies, resulting after taking the weighted sum of all layer models, as the *M* voxels. The regression procedure was the same as that used in the cnn2vox model. Response predictions from the cnn2vox and vox2vox models were thereafter combined as a weighted sum. The weight was determined by the relative accuracy of each prediction for each voxel.

#### 2.1.5. Vox2lab model

The vox2lab model estimates cognitive labels associated with movie scenes from predicted voxel responses. In this model, a series of z-scored cognitive labels at a time point *t*, denoted by L_(*t*)_(*t* = 1, …, *T*′) was regressed by a series of predicted responses to the scenes in the set of *N* voxels with hemodynamic delay, *k*, of 3, 4, and 5 seconds (Nishida & Nishimoto, 2018):

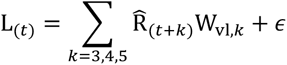

The regression procedure was the same as that used in the other models, except that the regularization parameters were determined separately for each dimension of label vectors. This model learns the association between predicted voxel responses (but not measured voxel responses) and cognitive labels.

### 2.2. Datasets

[Dum]

### 2.3. Movies

Two sets of movies were used in these experiments and analyses. One included 368 Japanese ad movies broadcasted on the web between 2015 and 2018 (web ad movie set). The other included 420 Japanese ad movies broadcasted on TV between 2011 and 2017 (TV ad movie set). The movies were all unique, included a wide variety of product categories (Table 1), and had the same resolution (1280 pixel × 720 pixel) and frame rate (30 Hz). The lengths of the movies were either 15 or 30 seconds. The movies were accompanied by PCM sound with a sampling rate of 44100 Hz and were normalized so that they had the same RMS level. To create movie stimuli for these experiments, the original movies in each set were sequentially concatenated in a pseudo-random order. Each movie set had a total length of 8400 seconds and was divided into 7200 seconds and 1200 seconds to collect voxel responses for the training (training dataset) and test (test dataset), respectively, of all the three models.

**Table 1.** The number of ad movies by product/service category.

| Categories | Web ad movie set | TV ad movie set |
| --- | --- | --- |
| Electronic & Precision | 4 | 9 |
| Audiovisual | 5 | 1 |
| Appliance | 16 | 5 |
| Car | 31 | 33 |
| Food & Confectionery | 7 | 69 |
| Beverage & Alcoholic drink | 20 | 38 |
| Medical & Health | 35 | 21 |
| Cosmetics | 49 | 6 |
| Sundries & Home equipment | 10 | 45 |
| Garment/apparel | 9 | 11 |
| Entertainment | 42 | 43 |
| Media & Education | 41 | 16 |
| Distribution & Retailer | 12 | 20 |
| Communication & Service | 35 | 51 |
| House & Construction | 9 | 12 |
| Finance | 9 | 20 |
| Enterprise, Public service, & Others | 34 | 20 |

### 2.4. Participants

Participants included Japanese individuals assigned to the fMRI experiments, forty (15 females; aged [mean ± SD] = 26.6 ± 9.0) with the web ad movie set and twenty-eight (12 females; age [mean ± SD] = 26.4 ± 7.6) with the TV ad movie set, with 16 participants overlapping between the two experiments. The experimental protocol was approved by the ethics and safety committees of NICT. Written informed consent was obtained from all participants.

### 2.5. MRI experiments

A 3T Siemens MAGNETOM Prisma scanner was used with a 32-channel Siemens volume coil and a multiband gradient echo-EPI sequence (Moeller et al., 2010) (TR = 1000 ms; TE = 30 ms; flip angle = 60 deg; voxel size = 2 × 2 × 2 mm^3^; matrix size = 96 × 96; the number of axial slices = 72; multiband factor = 6). The field of view covered the entire cortex. Using a T1-weighted MPRAGE sequence on the same 3T scanner, anatomical data were also collected (TR = 2530 ms; TE = 3.26ms; flip angle = 9 deg; voxel size = 1 × 1 × 1 mm^3^; matrix size = 256 × 256; the number of axial slices = 208).

In these experiments, the participants viewed the movie stimuli on a projector screen inside the scanner (spanning 27.9 deg × 15.5 deg of visual angle) and listened to the accompanying sounds through MR-compatible headphones. The participants were given no explicit task. The fMRI response data for both the web and TV ad movie sets were collected from individual participants in three separate recording sessions over 3 days.

For each stimulus set, 14 non-overlapping movie clips of 610-s length were obtained. The individual movie clips were displayed in separate scans. The initial 10-s part of each clip was a dummy to discard hemodynamic transients caused by clip onset, and the fMRI responses collected during the 10-s dummy movie were not used for the modeling. Twelve of the 14 clips were presented once to collect fMRI responses for model training (7200 s in total). The other two clips were presented four times each in four separate scans. The fMRI responses to these clips were averaged across four scans and used for model test (1200 s in total).

### 2.6. MRI data pre-processing

Motion correction in each functional scan was performed using SPM8. For each participant, all volumes were aligned to the first image taken in the first functional run. Low-frequency fMRI response drift was detected by using a median filter with a 120-s window and subtracting the filtered copy from the signal. Then, the response in each voxel was transformed into z-scoring responses to have zero-mean and unit variance. FreeSurfer (Dale et al., 1999; Fischl et al., 1999) was used to identify cortical surfaces from anatomical data and register them to the voxels of functional data. All voxels identified within the whole cerebral cortex for each participant were used for the modeling. In addition, cortical voxels were anatomically segmented into 358 regions based on the HCP-MMP1 atlas (Glasser et al., 2016) to show the localization of model performance on predicting voxel response.

### 2.7. Cognitive labels

We used five distinct categories of cognitive labels associated with movie scenes to evaluate the validity of predicted-response decoding (see also Table A.1).

#### 2.7.1. Scene descriptions

Manual scene descriptions using Japanese language were given for every 1-s movie scene of each of the web and TV ad movie sets. The annotators were 109 native Japanese speakers (79 females, age 19–62 years), who were not the fMRI participants. They were instructed to describe each scene (the middle frame of each 1-s clip) using more than 50 Japanese characters. Multiple annotators (5 and 12–14 annotators for web and TV ad movie sets, respectively) were randomly assigned for each scene to reduce the potential effect of personal bias. The descriptions contain a variety of expressions reflecting not only their objective perceptions but also their subjective perceptions (e.g., feeling; see Nishida et al., 2020 for examples of the descriptions). To evaluate the scene descriptions quantitatively, the descriptions were transformed into vectors of word2vec (Mikolov et al., 2013) in the same way as in previous work (Nishida et al., 2020, 2021; Nishida & Nishimoto, 2018). Individual words in each description were projected into a pretrained word2vec vector space. Then, for each scene, the word vectors obtained from all descriptions of the same scene were averaged. This procedure yielded one 100-dimensional vector for each 1-s scene. Thus, one label set was assigned to each movie set.

#### 2.7.2. Impression ratings

Manual ratings given for every 2-s movie scene on 30 different impression labels (e.g., “beautiful”; Table A.1) were collected from 44 human annotators (34 females, age 20–61 years), who were different from the fMRI participants, except for 6 annotators for the TV ad movie set. The annotators sequentially watched separate 2-s clips of the movies with sound and evaluated each label on a 5-point scale. To keep the motivation of the annotators, only 15 labels were assigned to each annotator in single clip evaluation. Every label was evaluated from 9 different annotators for each scene of the web ad movie set and from 12 different annotators for each scene of the TV ad movie set. The mean impression rating for each label and each scene was obtained by averaging ratings over these annotators. Then the mean ratings were oversampled to obtain time series of rating labels in every 1-s scene. Thus, 30 labels were assigned to each movie set.

#### 2.7.3. Ad effectiveness indices

Two types of mass behavior indices were collected from the Internet. One is click-through rate, or the fraction of viewers who clicked the frame of a movie and jumped to a linked web page. The other is view completion rate, or the fraction of viewers who continued to watch an ad movie until a specific time point of the movie (25%, 50%, 75%, or 100% from the start) without choosing the skip option. Hence, each movie had one index of click-through rate and four indices of view completion rate. These indices were computed from 3783 to 12268028 (mean = 632516) unique accesses for each movie. The values were oversampled to obtain a time series of indices in every 1-s scene. Thus, a total of five labels were assigned to the web ad movie set.

#### 2.7.4. Ad preference votes

Reputation surveys of TV ads for commercial purposes were conducted using questionnaires to large-scale testers. Each tester was asked to freely recall a small number of her/his favorite TV ads from among the ads recently broadcasted. The total number of recalls of an ad was regarded as a preference vote. In addition, the questionnaires also included 15 subordinate items that asked why the ads were favorable to the viewer (e.g., because it was “humorous”), and three additional items indexed how effective the ads were for the viewer’s using and purchasing the products (e.g., “purchase intention”; Table A.1). To eliminate the bias of preference due to frequent exposure on TV, the number of votes for each item of an ad was divided by its gross rating point (GRP); GRP is an index of how many people see an ad over a particular period, and calculated by the audience ratio multiplied by the number of times the ad is broadcasted during the period. The values were oversampled to obtain a time series of votes and items in every 1-s scene Since these values are distributed in a similar form of a gamma distribution, the logarithm of the data was taken. Thus, a total of 19 label sets were assigned to the TV ad movie set.

#### 2.7.5. Subjective preference ratings

Manual preference ratings given for each TV ad movie were collected from 14 of the fMRI participants on separate days after fMRI experiments. While the participants sequentially watched each movie outside the MRI scanner, they rated their own preference for the movie on a 9-point scale, from most unlikable to most likable. To keep the motivation of the participants, the ratings were conducted in separate seven blocks (600 s per block) with intervals on each of two days. The ratings were oversampled to obtain a time series of ratings in every 1-s scene. This resulted in one label set assigned to the TV ad movie set.

### 2.8. Model construction

For the predicted-response decoding, the three models for each participant were constructed using the paired data obtained for the web or TV ad movie set, i.e., the participant’s fMRI data and cognitive labels in the training data. After the cnn2vox and vox2vox models were trained using movies and the measured voxel responses to them, the vox2lab model for each label was trained using the predicted voxel responses to the same movies and the label associated with them. Then, using the test data, a time series of each label was estimated using these three models.

To measure decoding accuracy for each label, the Pearson correlation coefficient between this estimated time series and a time series of the true label was calculated for each participant. Note that since the label of scene description used multidimensional vectors, the decoding accuracy for this label was evaluated by calculating the correlation coefficient at each time point and averaging it over all time points. To evaluate whether the decoding accuracy for a given label was significantly greater than zero was evaluated with the Wilcoxon signed-rank test with correction for multiple comparisons using the false discovery rate (FDR), drawing on each participant’s accuracy as a data sample (P < 0.05).

For the comparison with predicted-response decoding, we also constructed each participant’s model for measured-response decoding, in which cognitive labels were directly estimated by decoding measured voxel responses to movies. The form of this model was the same as in the vox2lab model of the predicted-response decoding, except that measured voxel responses were used instead of predicted responses. The regression procedure and the significance test were the same as in the predicted-response decoding.

### 2.9. Individual difference

We introduced a measure called individual-difference reflection (IDR) to evaluate the similarity between individual differences obtained from predicted-response decoding and those from measured-response decoding (Fig. 2). First, we assessed the individual differences in cognitive labels estimated with each decoding method. This was done by evaluating the participant-pair dissimilarity in the time series of estimated labels for the test data. This dissimilarity was determined using the Pearson correlation distance (i.e., 1 − Pearson correlation coefficient) between all possible participant pairs for each movie set. We performed this separately for both predicted- and measured-response decoding.

**Fig. 2.**
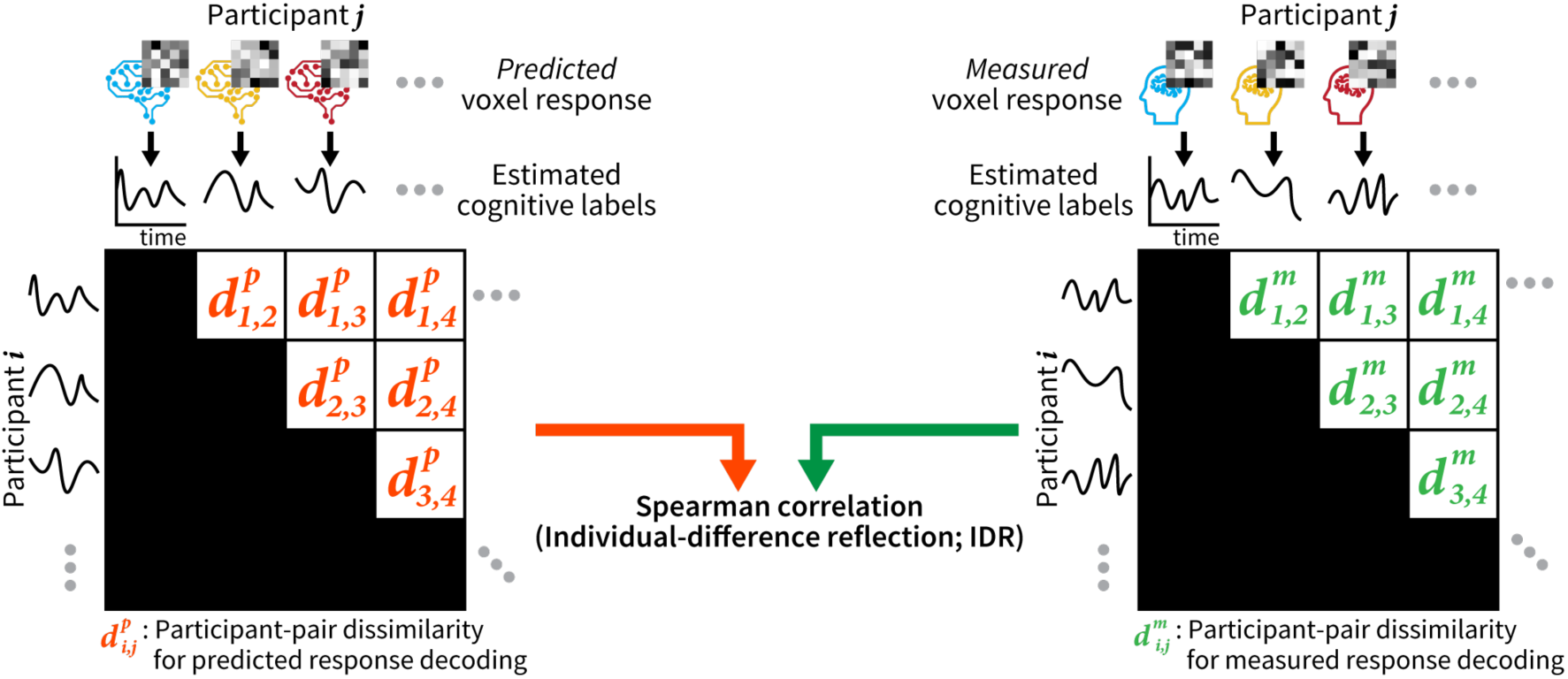
Individual-difference reflection (IDR). The IDR measures how accurately the predicted-response decoding framework captures individual differences in mental information. First, for each pair of participants and each cognitive label, we calculated the participant-pair dissimilarity (1 – Pearson correlation coefficient) between the time series of the same label estimated for the two participants by predicted-response decoding (left); and did the same for measured-response decoding (right). Next, for each cognitive label, the IDR was evaluated by the Spearman correlation between the participant-pair dissimilarities obtained with the predicted-response decoding and those with the measured-response decoding.

Then, for each label, we calculated the IDR. It is defined as the Spearman correlation coefficient between the participant-pair dissimilarities derived from predicted-response decoding and those from the measured-response decoding for the same label. Note that we opted for Spearman correlation, as it is previously recommended for computing the correlation of dissimilarity measures (Nili et al., 2014). However, we observed comparable results when using Pearson correlation (see Results). When the IDR (Spearman correlation) for a given label was significantly larger than 0 (P < 0.05, FDR corrected), it indicated that the predicted-response decoding reflected the individual differences in mental information for that label.

## 3. Results

### 3.1. Model performance

To validate the models constructed for the predicted-response decoding, we first evaluated the model performance at the population level in terms of voxel-response prediction and decoding. The accuracy of voxel-responses prediction (prediction accuracy) in the cnn2vox and vox2vox models was evaluated by their similarity to measured voxel responses, as quantified by their Pearson correlation (Fig. 3). We found that the visual CNN-based models accurately predicted voxel responses in visual regions, such as occipital and posterior temporal cortical areas. In contrast, the auditory CNN-based models predicted voxel responses in auditory regions, such as anterior temporal cortical areas. Theses localized patterns of the prediction accuracy indicate that the models based on the visual and auditory CNNs selectively predicted voxel responses in the visual and auditory regions of the cortex, respectively. This result is consistent with previous findings in brain-response modeling studies using CNNs (Güçlü & van Gerven, 2015, 2017; Kell et al., 2018).

**Fig. 3.**
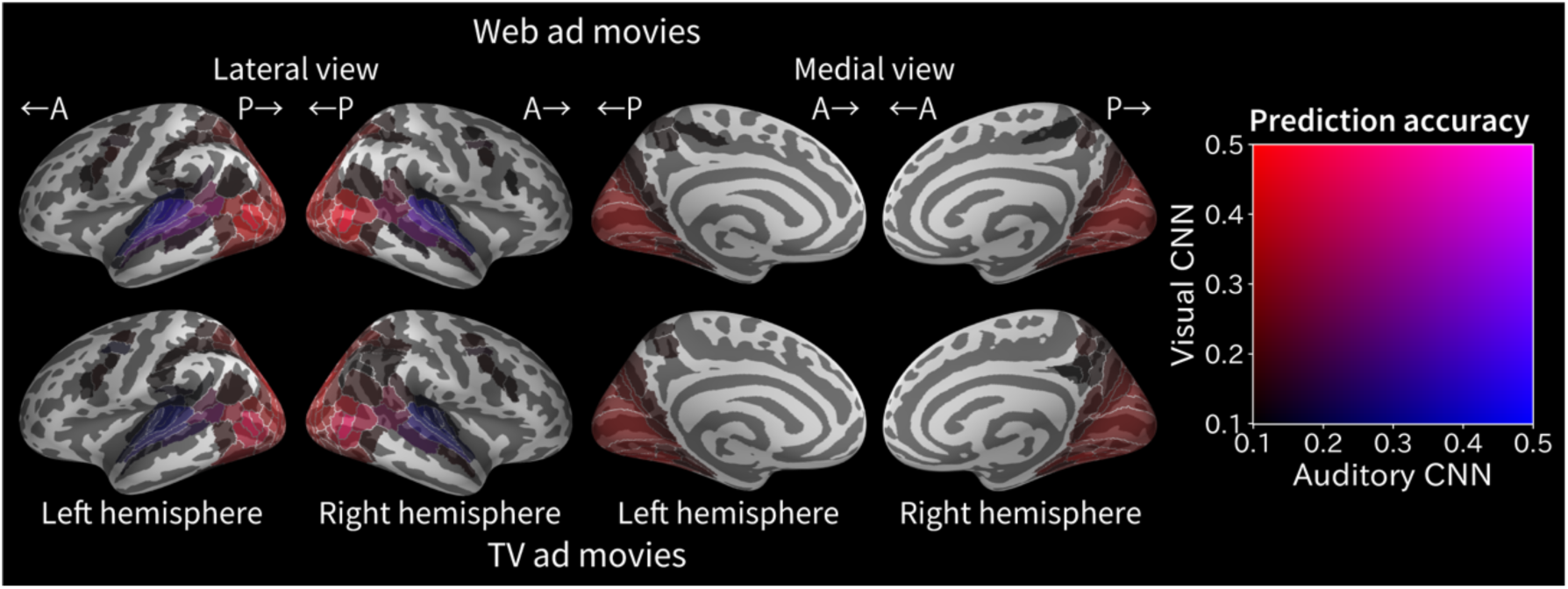
Localization of cortical regions in which the response is predicted by cnn2vox and vox2vox models. Participant-average (top: web ad movie set, n = 40; bottom: TV ad movie set, n = 28) prediction accuracy is shown within each of the 358 cortical regions segmented based on the HCP-MMP1 atlas. Color indicates the mean prediction accuracy within each of the regions, as denoted by the colormap (right). Accuracy less than 0.1 is not shown. The regions predicted preferentially by the visual CNN-based models and the regions predicted preferentially by the auditory CNN-based models appear in distinct colors (red and blue, respectively). A, anterior; P, posterior.

The decoding accuracy of the vox2lab model was evaluated by the Pearson correlation between decoded and true cognitive labels. The accuracy was significantly higher than 0 for all 87 cognitive labels (range, 0.11–0.82; Wilcoxon signed-rank test, P < 0.0005, FDR corrected) and strongly correlated with that of measured-response decoding across all labels (Pearson r = 0.69, P < 0.0001; Fig. 4). Furthermore, the accuracy was rather higher for predicted-response decoding than for measured-response decoding (Wilcoxon signed-rank test, P < 0.0001), as reported previously (Nishida et al., 2020). Thus, at the population level, these models effectively estimate mental information.

**Fig. 4.**
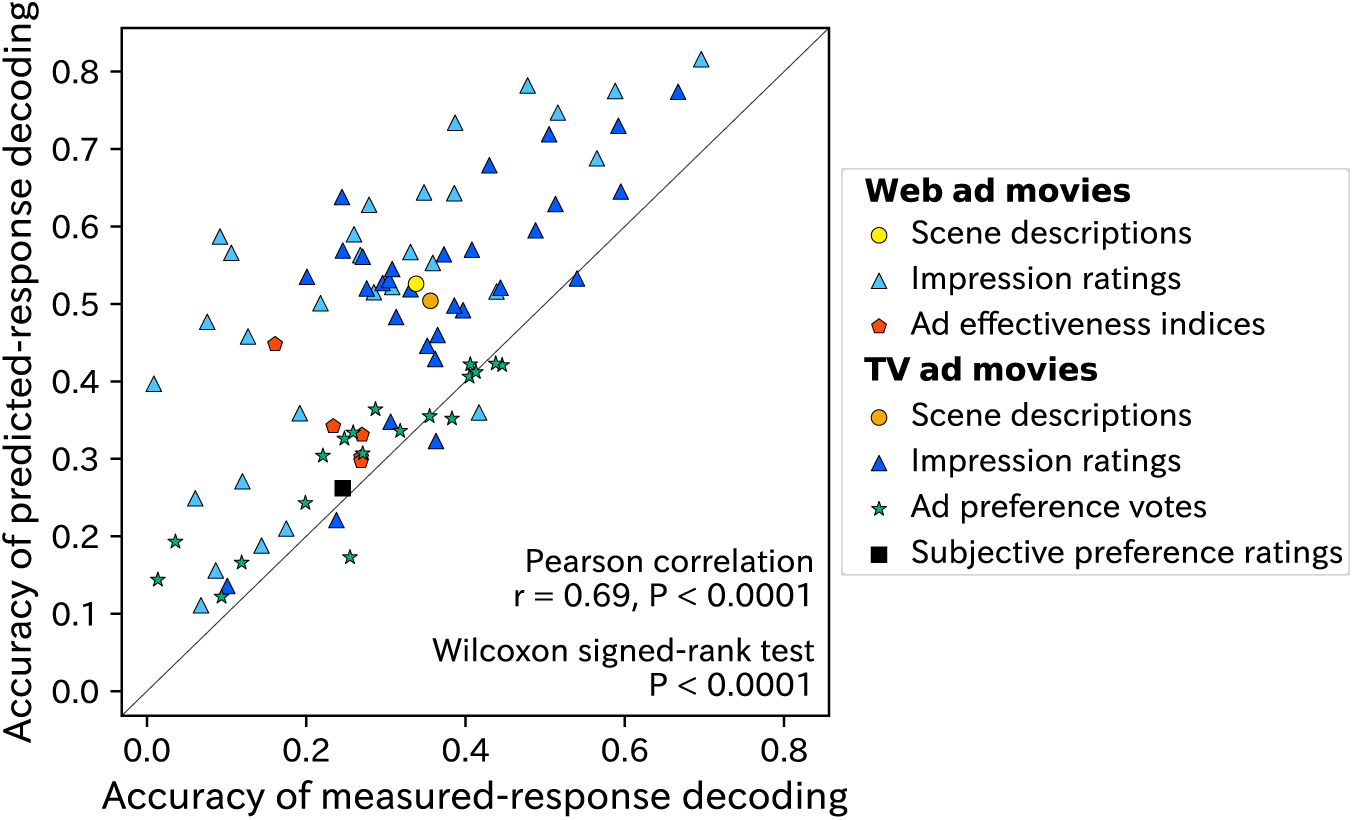
Label-estimation accuracy compared for predicted- and measured-response decoding. The participant-average accuracy for predicted-response decoding (y axis) is plotted against that of measured-response decoding (x axis). Each symbol denotes accuracy for one cognitive label.

### 3.2. Significant IDR for almost all cognitive labels

We next examined how well predicted-response decoding captures the individual differences in mental information derived from the measured-response decoding. Among all 87 cognitive labels, the highest value of the IDR was observed for the label of scene descriptions for the web ad movie set (0.74; Fig. 5A). The estimation of this label was derived from the paired data of predicted voxel response and cognitive label, each of which was derived from different persons. However, even when using the pair data derived from the same person (i.e., when estimating the label of subjective preference ratings), we obtained a significant value of the IDR (P < 0.0005; Fig. 5B).

**Fig. 5.**
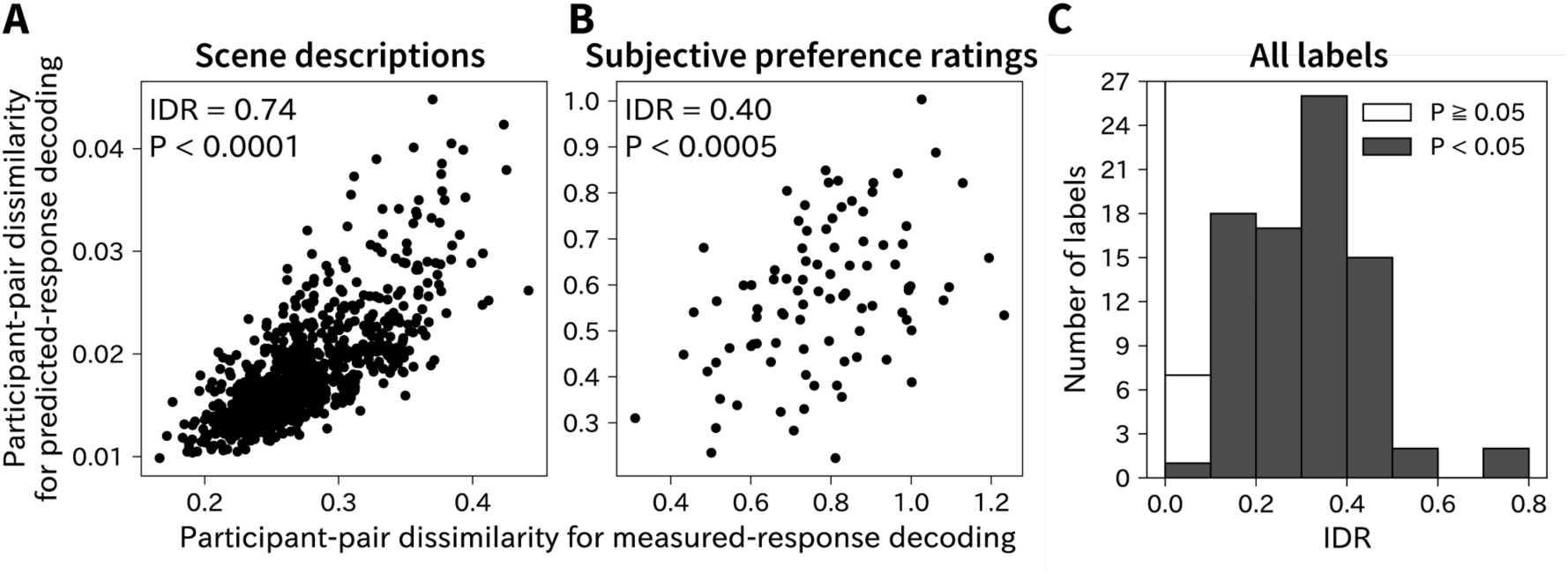
IDR for two example labels and for all 87 labels. (A, B) Two example labels showing significant IDR: the label of scene descriptions for the web ad movie set (A) and the label of subjective preference ratings for the TV ad movie set (B). The participant-pair dissimilarity of the label obtained with predicted-response decoding (y axis) are plotted against those obtained with the measured-response decoding (x axis). Each dot represents one pair of participants. (C) The distribution of IDR for all 87 labels. Filled bars indicate significant IDR (Spearman correlation, P < 0.05, FDR corrected).

Of all the 87 cognitive labels decoded in this study, we observed significant IDR for 81 labels (Fig. 5C and Table A.2; P < 0.05 with correction for multiple comparisons using FDR). A similar result was observed even when a different measure (i.e., Pearson correlation instead of Spearman correlation) was used to calculate the IDR (Fig. A.1). Thus, these results suggest that predicted-response decoding successfully captures the individual differences in mental information for almost all the cognitive labels used across different datasets.

Are individual differences in mental information captured by predicted-response decoding associated with individual differences in subjective perceptions? We examined this by testing the correlation (IDR) between subjective preference ratings estimated using predicted-response decoding and those given manually by participants, resulting in a significant IDR (0.54, P < 0.0001; Fig. 6). These findings suggest that predicted-response decoding can capture individual differences not only in mental information derived from measured-response decoding but also in subjective perceptions.

**Fig. 6.**
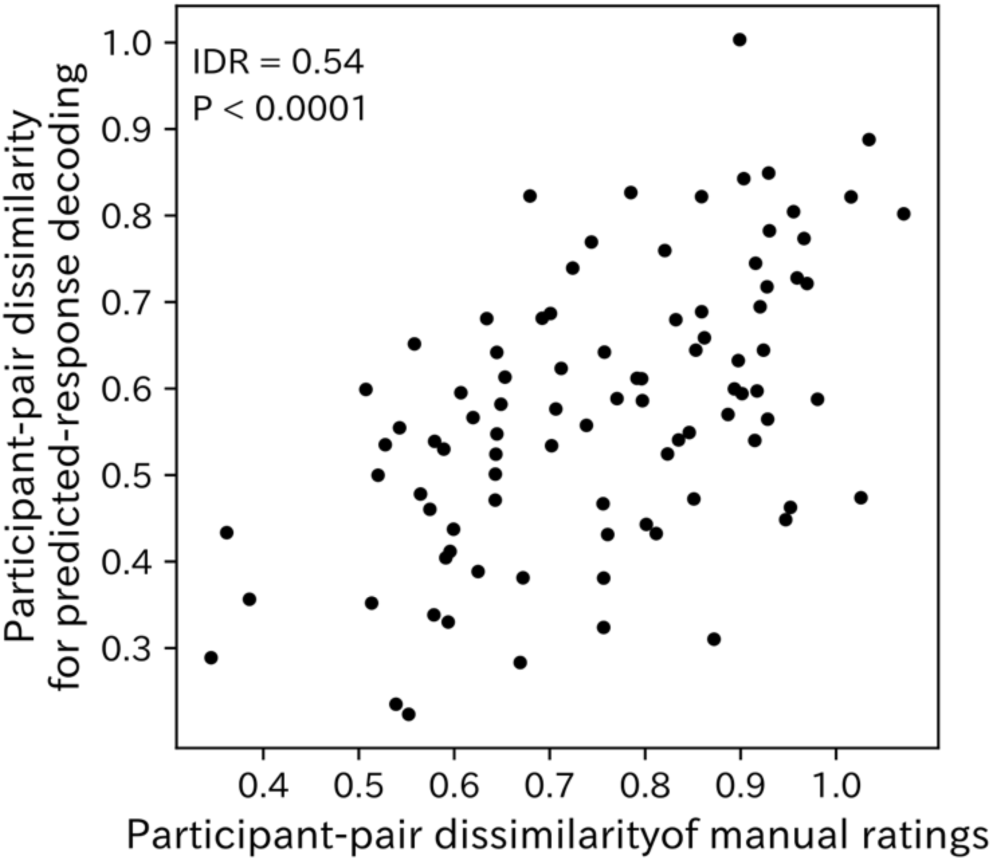
IDR for subjective preference ratings. The participant-pair dissimilarity of subjective preference ratings estimated with predicted-response decoding (y axis) are plotted against those of manual ratings (x axis). Each dot represents the participant-pair dissimilarity of one pair.

### 3.3. Relationship between IDR and decoding performance

The IDR, however, varied across cognitive labels, raising the question of what factor determines the variation of the IDR. We tested the hypothesis that the magnitude of the IDR for a given label is affected by the accuracy of the predicted-response decoding for that label (Fig. 7). A significant correlation between the IDR and decoding accuracy (Pearson r = 0.47, P < 0.0001) supports this hypothesis. This suggests that the more accurately we decode a cognitive label using the predicted-response decoding, the more effectively the decoded contents reflect the individual differences in mental information for the label.

**Fig. 7.**
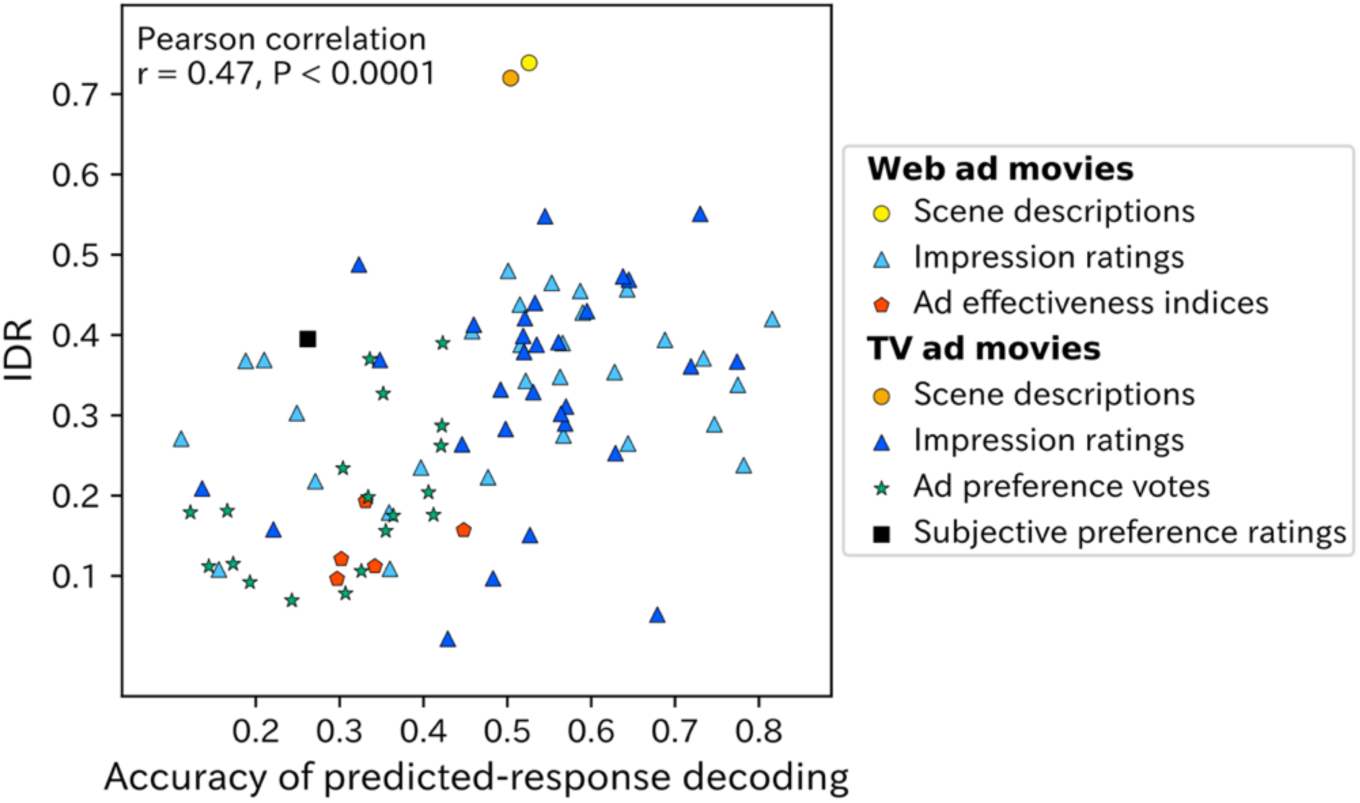
Correlation between IDR and the accuracy of predicted-response decoding. IDR for each cognitive label (y axis) is plotted against the participant-average accuracy of predicted-response decoding for that label (x axis). Each symbol denotes one cognitive label.

## 4. Discussion

We introduced a novel brain decoding framework, called predicted-response decoding, to estimate the individual differences in mental information evoked by natural movie scenes with minimal fMRI measurement (Fig. 1). We found that this decoding framework successfully captured the patterns of individual differences in mental information, when compared with those derived from conventional measured-response decoding, for 81/87 (93.1%) of decoded cognitive labels associated with natural movie scenes (Fig. 5). This result suggests that the predicted-response decoding framework can be used to estimate personal mental information evoked by natural scenes.

The novelty of this study lies in the successful decoding of individual differences in mental information from predicted brain responses instead of measured ones. While recent advancements in brain decoding have achieved significant progress in tasks like image reconstruction (Takagi & Nishimoto, 2023) and in language generation (Tang et al., 2023) using brain responses, all existing approaches have relied on decoding from measured brain responses. This study, however, takes an entirely distinct approach by decoding from predicted brain responses—a novel strategy with the potential to considerably facilitate real-world applications of brain decoding by virtue of substantial reductions of measurement costs.

The methodology utilized in this study is based on the technique we have previously proposed (Nishida et al., 2020); however, the previous study employed this technique as a means of transfer learning. The decoding of mental information, encompassing individual differences, from predicted brain responses marks a completely different application of this technique. Hence, the unique contribution of this study lies in introducing a completely novel application of an existing methodology. Furthermore, the rigorous validation of this application using multiple datasets and diverse estimation tasks also serves as a noteworthy contribution of this study.

To reduce the costs and constraints of fMRI measurements, previous studies have mainly proposed methods for the inter-individual alignment of the representational space in the brain (Conroy et al., 2013; Guntupalli et al., 2016; J. V. Haxby et al., 2011; Yamada et al., 2015). These methods incur small measuring costs and have great capability for the inter-individual generalization of brain decoders. However, our proposed framework has unique advantages against these methods. First, our framework requires no fMRI measurement to estimate mental information induced by *novel* audiovisual inputs. Second, our framework can construct new decoders without any additional fMRI measurements because the decoders are trained using predicted (not measured) brain responses. In addition, our framework is compatible with the inter-individual alignment methods. The models for brain-response prediction in our framework may be generalized across individuals via inter-individual alignment. In fact, such combination use could further reduce the fMRI-associated costs and constraints for brain decoding.

We evaluated the predicted-response decoding framework using moderate performance CNNs (i.e., VGG-16 and SoundNet) for extracting features from movies and using linear regression for mapping brain responses with movie features or cognitive labels, according to the settings in a previous study (Nishida et al., 2020). Although the framework successfully captures the individual differences in mental information even for these settings, model performance, as measured by the individual-difference reflection, may be improved by employing higher-performance CNNs (Schrimpf et al., 2020) and/or more sophisticated nonlinear regression, such as regression with deep neural networks (Khosla et al., 2021). It is important to note that our framework essentially allows us to apply, as plug-ins, any methods to perform feature extraction and regression. Therefore, future investigations should address optimization of component methods to be used for predicted-response decoding of individual differences.

By utilizing predicted brain responses in place of measured ones, the costs associated with brain measurements are drastically reduced. This reduction in cost brings about a pivotal contribution in the field of neuromarketing, one of the promising applications of brain decoding (He et al., 2021; Plassmann et al., 2015). Neuromarketing leverages mental information decoded from brain responses to shape marketing strategies for services and products. However, the substantial costs tied to brain measurements have been a major obstacle to the progression of neuromarketing. The introduction of the predicted-response decoding framework offers a potential solution to this challenge.

## 5. Conclusion

Recent fMRI-based brain decoding has achieved remarkable success in recovering rich mental information from brain responses to natural scenes. This holds promising potential for real-world applications. However, the high costs of the required fMRI measurements limit its widespread use. To address this, we proposed a novel decoding framework where mental information evoked by natural scenes is recovered from predicted brain responses instead of measured ones. We found that this predicted-response decoding successfully captures individual differences in mental information derived from measured-response decoding across various tasks for estimating cognitive labels, even when these labels are associated with subjective perceptions. Thus, predicted-response decoding provides a cost-effective, versatile tool for brain decoding to estimate personal perceptions evoked by natural scenes, greatly enhancing real-world applications of brain decoding.

## Appendix

**Fig. A.1.**
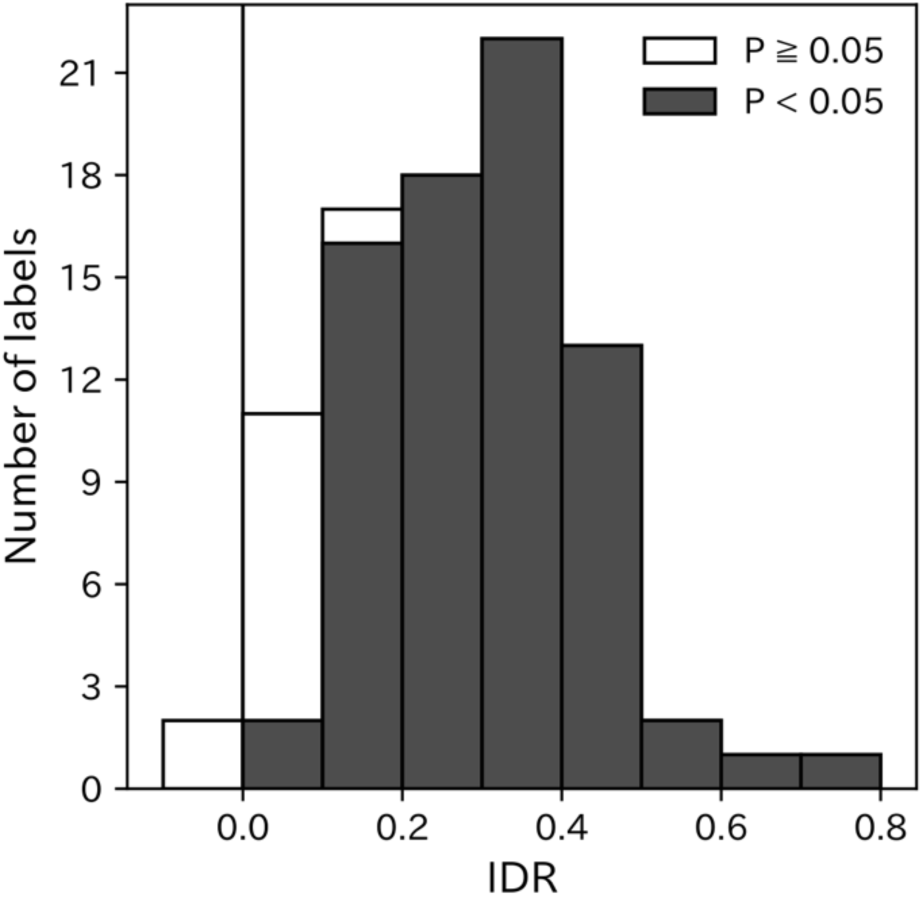
IDR for all cognitive labels when Pearson correlation was used instead of Spearman correlation. IDR was significant (gray bars; P < 0.05, FDR corrected) for 75 of 87 labels.

**Table A.1.**
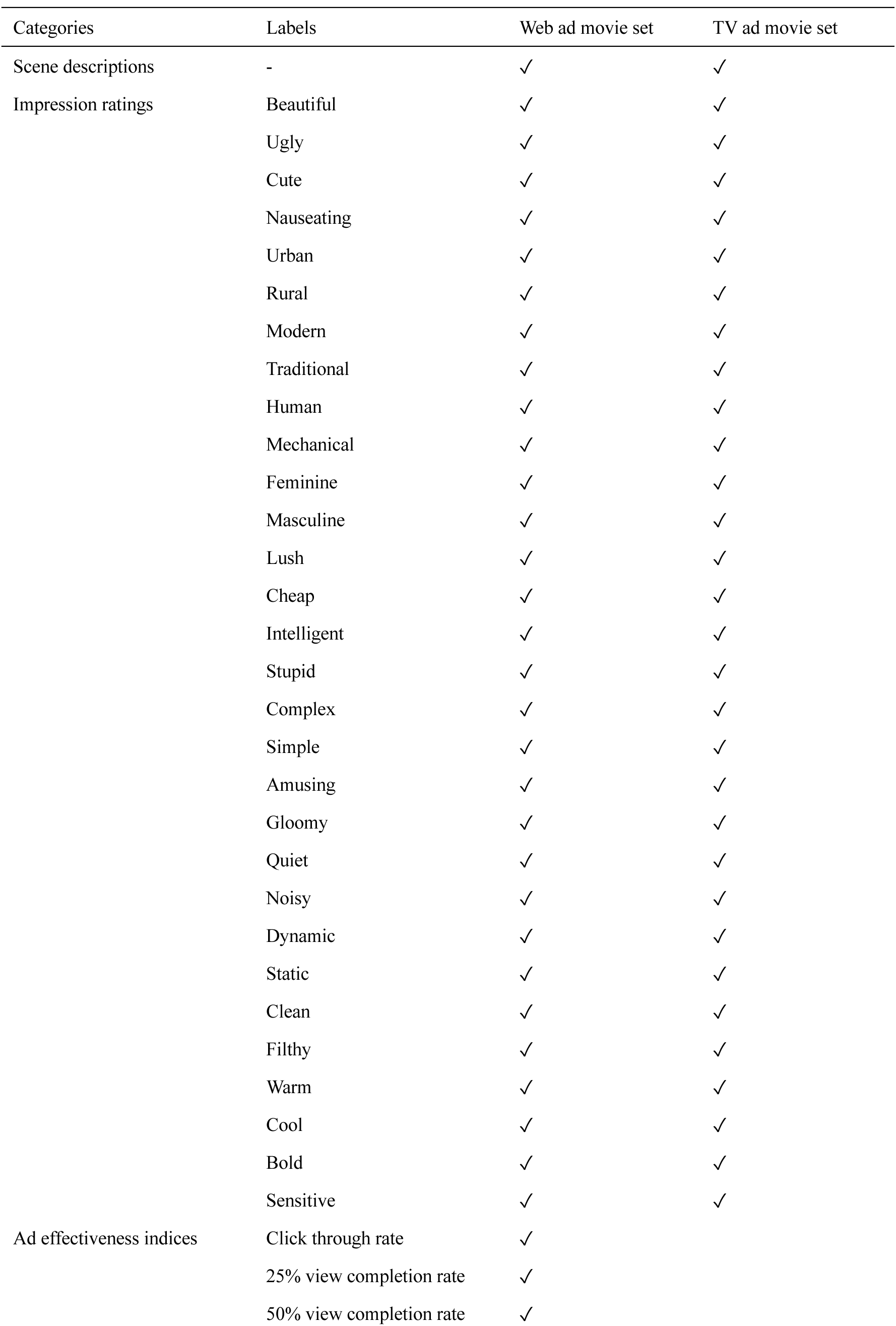
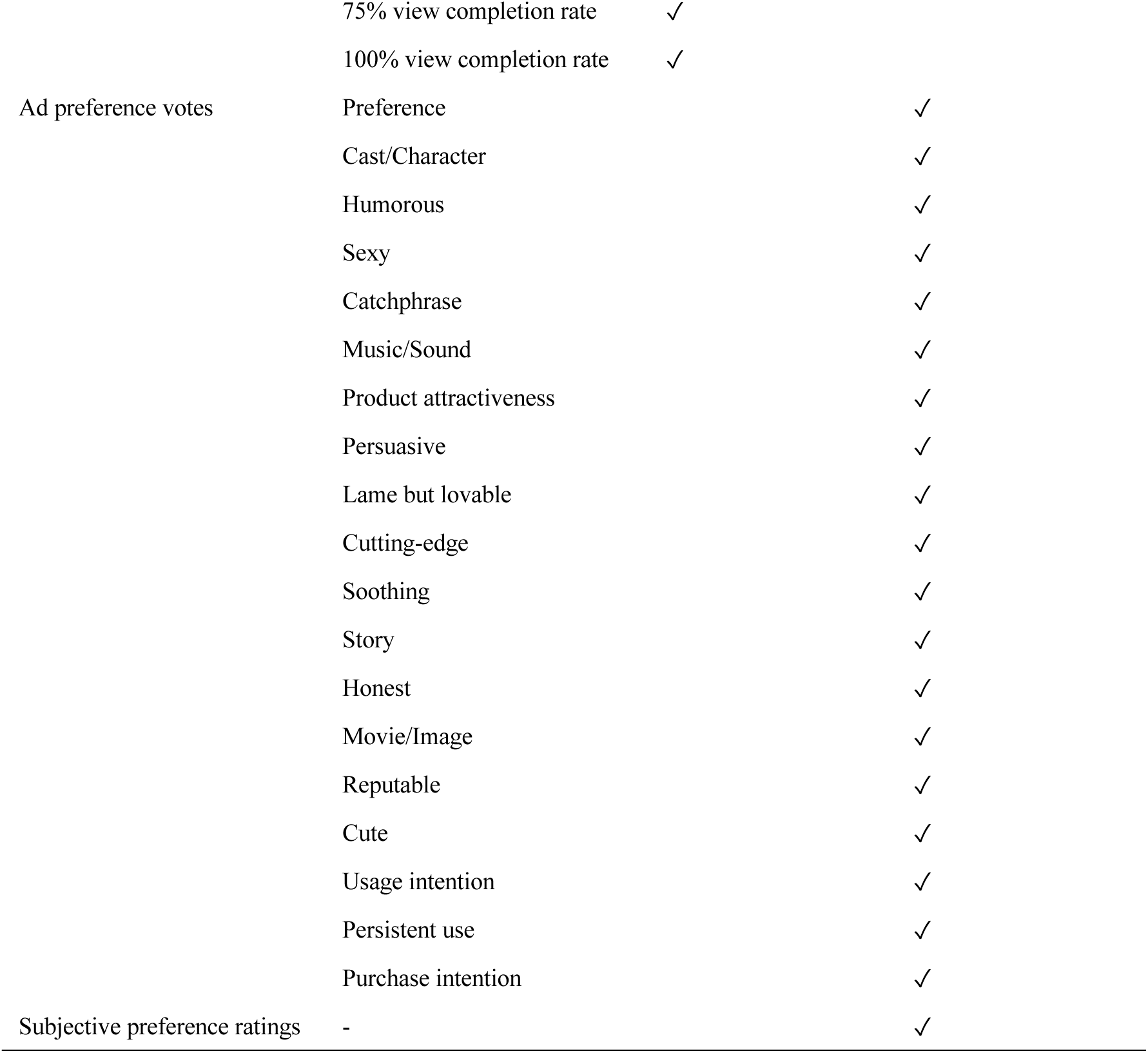
Cognitive labels associated with each movie set.

**Table A.2.** IDR for cognitive labels. *P < 0.05, **P < 0.01, ***P < 0.0001, all FDR corrected.

| Category | Cognitive label | IDR |  |
| --- | --- | --- | --- |
|  |  | Web ad movie set | TV ad movie set |
| Scene descriptions | - | 0.739*** | 0.720*** |
| Impression ratings | Beautiful | 0.354*** | 0.332*** |
|  | Ugly | 0.271*** | 0.158** |
|  | Cute | 0.457*** | 0.399*** |
|  | Nauseating | 0.303*** | 0.369*** |
|  | Urban | 0.455*** | 0.388*** |
|  | Rural | 0.405*** | 0.473*** |
|  | Modern | 0.223*** | 0.290*** |
|  | Traditional | 0.218*** | 0.097 |
|  | Human | 0.394*** | 0.469*** |
|  | Mechanical | 0.465*** | 0.253*** |
|  | Feminine | 0.371*** | 0.302*** |
|  | Masculine | 0.265*** | 0.151** |
|  | Lush | 0.275*** | 0.413*** |
|  | Cheap | 0.368*** | 0.283*** |
|  | Intelligent | 0.369*** | 0.421*** |
|  | Stupid | 0.109** | 0.440*** |
|  | Complex | 0.179*** | 0.488*** |
|  | Simple | 0.338*** | 0.311*** |
|  | Amusing | 0.343*** | 0.361*** |
|  | Gloomy | 0.235*** | 0.022 |
|  | Quiet | 0.238*** | 0.551*** |
|  | Noisy | 0.289*** | 0.052 |
|  | Dynamic | 0.420*** | 0.367*** |
|  | Static | 0.348*** | 0.329*** |
|  | Clean | 0.428*** | 0.379*** |
|  | Filthy | 0.108** | 0.209*** |
|  | Warm | 0.480*** | 0.264*** |
|  | Cool | 0.390*** | 0.391*** |
|  | Bold | 0.388*** | 0.548*** |
|  | Sensitive | 0.438*** | 0.430*** |
| Ad effectiveness indices | Click through rate | 0.157*** |  |
|  | 25% view completion rate | 0.121** |  |
|  | 50% view completion rate | 0.193*** |  |
|  | 75% view completion rate | 0.096** |  |
|  | 100% view completion rate | 0.112** |  |
| Ad preference votes | Preference |  | 0.204*** |
|  | Cast/Character |  | 0.262*** |
|  | Humorous |  | 0.287*** |
|  | Sexy |  | 0.179** |
|  | Catchphrase |  | 0.115* |
|  | Music/Sound |  | 0.390*** |
|  | Product attractiveness |  | 0.078 |
|  | Persuasive |  | 0.112* |
|  | Lame but lovable |  | 0.106* |
|  | Cutting-edge |  | 0.181** |
|  | Soothing |  | 0.175** |
|  | Story |  | 0.176** |
|  | Honest |  | 0.092 |
|  | Movie/Image |  | 0.370*** |
|  | Reputable |  | 0.070 |
|  | Cute |  | 0.234*** |
|  | Usage intention |  | 0.156** |
|  | Persistent use |  | 0.198** |
|  | Purchase intention |  | 0.327*** |
| Subjective preference ratings | - |  | 0.395** |

## Declaration of generative AI in scientific writing

During the preparation of this work the authors used ChatGPT in order to improve readability and language. After using this tool/service, the authors reviewed and edited the content as needed and take full responsibility for the content of the publication.

## Acknowledgements

We thank Ms. Hitomi Koyama and Ms. Amane Tamai for their experimental support.

## Funding

This work was supported by the Japan Society for the Promotion of Science [grant number 21H03535]; the Japan Science and Technology Agency [grant numbers JPMJPR20C6 and JPMJSP2138]; NTT DATA Japan Corporation.

## Notes

### Competing Interest Statement

This study was funded by NTT Data Corp. NM is an employee of NTT Data Corp.

### Summary of Updates

We have made improvements throughout the content of the paper.

